# Expression profiling of immune-response genes common in *SARS-CoV-1* and *influenza A virus* disease, viz-a-viz neutropenia disorder, using integrated bioinformatics tools

**DOI:** 10.1101/2022.09.09.507303

**Authors:** Ayinde Olaniyi, Obasanmi Oluwadamilola, Alabi Oluwabunmi

## Abstract

Viral diseases have proven to be an existential threat to man due to its fatality and its consistent emergence and re-emergence, so the need for novel ideas in combating this menace. Advances in genetic studies has proven to be indispensable in this fight as knowledge of organismal genetic variability has been useful in therapy development, as well as in mounting defense against the treacherous infectious agents. However, there are still fallow grounds needing to be explored in this war which informed this study, with focus on expression profiling of similar immune-response-genes in SARS-COV-1 and influenza A virus (AIV) in respect to neutropenia (NP), using integrated bioinformatics techniques such as extraction of microarray dataset in the gene expression omnibus (GEO) database, generation of DEGs using GEO2R tool, construction of protein-protein interaction (PPI) network using string and Cytoscape software, Venn diagram analysis and then gene ontology (GO) and Kyoto encyclopedia of genes and genomes (KEGG) for enrichment and pathway analysis respectively. Ten genes which includes ELANE, ITGA2B, CXCR1, CSF3R, SPI1, MS4A3, MMP8, CEACAM8, RNASE3, and DEFA4 were identified, with ELANE gene moreover identified as a key gene, which might be responsible in the regulation of immune response during viral infection, based on its characteristic feature in the PPI network.

## 1. Introduction

The complex interplay between the host gene and genetic components of polygenic microorganisms such as viruses has a significant contributory role to the susceptibility to infectious diseases [1], hence the focus of research towards the manufacture of several measures and methods helpful in untangling this complexity in the fight against this menace. However, the advances in genome sequencing and genomic studies has provided invaluable knowledge in enumerating the variability in genetic construct of different organisms, vital to raising appropriate countermeasure against invading infectious and non-infectious diseases [2]. Genes are known to carry out multiple biological functions [3], however the primary biological function of a gene can also be verified based on the expression level in varying conditions which may allow for the possible manipulation of theses gene, important in disease control and treatment.

Following the aforementioned, the comparative studies of differentially expressed genes reveal the degree of similarity that exists between different diseases, which proves invaluable to understanding and appreciating disease biology [4]. Riding on this premise, and the due to absence of this study in literature, this study seeks to investigate the immune response genes common in both SARS-COV-1 and influenza A virus (IAV) in comparison to the differentially expressed genes of neutropenia (NP), using the miRNA dataset; miRNAs are small non-coding RNAs containing ~22 nucleotides, which serve important functions in RNA silencing and post-transcriptional regulation of gene expression [5].

SARS-COV-1 and IAV belongs to the severe acute respiratory tract infections(SARI) known as the fourth most common cause of death worldwide [6].The SARS-COV-1 belong to the species SARS-related coronavirus (subgenus sarbecovirus, genus betacoronavirus) with approximately 80% genome sequence similarity with the latest SARS-COV-2 [7], responsible for the SARS outbreak in 2002–2003, with more than 8000 infections and more than 700 deaths worldwide [8]. The influenza A virus (IAV) is the most prevalent agent among SARI viruses as the first breakout of IAV in 1918 reportedly claimed around 50 million lives worldwide, with seasonal yearly outbreaks claiming around 650 thousand lives [6]. NP on the other hand has been reported to be characterized significantly by lower-than-normal absolute neutrophil count, with the subjects at varying degrees’ risk of infection based on the cause and duration of this disorder. Individuals with neutrophil counts of 1.0–1.8 × 103/μl (1.0–1.8 × 109/liter) are at little risk, while between 0.5 × 103 and 1.0 × 103/μl (0.5–1.0 × 109/liter) neutrophils with a moderate risk. Individuals with neutrophil counts of less than 0.5 × 103/μl (0.5 × 109/liter) are at substantially greater risk [9].

The background knowledge of the central role played by neutrophil in innate immunity [10] informs the choice of Neutropenia (NP);which is characterized by low neutrophil content, in this study. During an infection, neutrophil secretes neutrophil elastase; mutations of the ELANE gene which encodes neutrophil elastase causes cyclic and severe congenital neutropenia, which is a failure of neutrophils to mature [11]. The comparison of the top differentially expressed genes (DEGs) common in SARS-COV-1 and IAV with the DEGs of NP is therefore an important method put forward to check for the presence of these genes with the aim of investigating their contributory role in immune response.

## 2. Materials and methods

### 2.1. Retrieval of microarray data

The microarray gene expression dataset GSE1739, GSE6322, and GSE66597, were retrieved from the gene expression omnibus (GEO) database of NCBI (https://www.ncbi.nlm.nih.gov/geo/) for SARS-COV,IAV, and neutropenia respectively, using keywords SARS-COV, influenza A virus, and neutropenia.

**Table I.** Dataset information for microarray dataset obtained from GEO. 10 for SARS-COV patients and 4 normal samples as the control in dataset GSE1739, 2 IAV patients, and 2 normal samples as the control in dataset GSE6322. 9 neutropenia patients, and 9 normal Samples as the control in dataset GSE66597.

| DATASET | NO OF PATIENT | NO OF CONTROL |
| --- | --- | --- |
| GSE1739 | 10 | 4 |
| GSE6322 | 2 | 2 |
| GSE66597 | 9 | 9 |
| *Total* | 21 | 15 |

### 2.2. Identification of DEGs

Significant DEGs between test and control samples were analyzed using an online analysis tool GEO2R (https://www.ncbi.nlm.nih.gov/geo/geo2r/) for the dataset. The GEO2R tool is an interactive web tool for comparing two sets of data under the same experimental conditions and can analyze any geo series [12], it uses Bioconductor packages such as GEOQuery and limma for processing of data [13]. DEGs between the test blood samples and normal blood samples were screened with the following threshold criteria; P-value < 0.05 and a fold change value logFC ≥ 1.0.

### 2.3. Identification of common genes

The FunRich version 3.1.3 an online software was used for creating Venn Diagram through which we can get the common genes between our DEGs [14].

### 2.4. Integration of the protein-protein interaction (PPI) network

The PPI network of common genes was constructed using the online-based tool STRING (https://string-db.org/) for protein-protein interaction(PPI),with confidence score of ≥ 0.4 [15].

### 2.5. Identification of hub genes and source of interaction within genes

The constructed PPI network was visualized using desktop-based cytoscape software (3.7.1) (https://cytoscape.org/). Using the CytoHubba plugin of cytoscape, top 10 genes were identified defined by the MCC method; to finding the hub genes, maximal clique centrality (MCC) algorithm has been reported to be the most effective method [16]. In the same vein, the sources of interactions were identified using the style plug-in of the cytoscape.

### 2.6. Functional Annotation of common genes

The enrichment analyses were performed by uploading common genes into the online-based tool Enrichr (https://maayanlab.cloud/Enrichr/). Enrichr executes gene ontology (GO) which includes biological process, cellular component and molecular function of the gene, as well as executing the Kyoto Encyclopedia of Genes(KEGG) for pathway enrichment of the genes. GO is a common way to annotate genes, their products, and their sequences. while KEGG is a detailed data-base resource for the biological interpretation of genomic sequences and other high-throughput data.

### 2.7. microRNA network analysis

The common genes were submitted to the online software miRNet (https://www.mirnet.ca/) to find out the interaction between miRNAs and genes. Network between genes and miRNAs are generated using this software.

## 3. Results

### 3.2. Identification of DEGs

With the help of GEO2R tool, the datasets GSE1739, GSE6322, and GSE66597 analysis returned 407,1649, and 246 DEGs respectively with a set cut-off criteria | logFC | ≥1.0 and P < 0.05.

**Figure 1.**
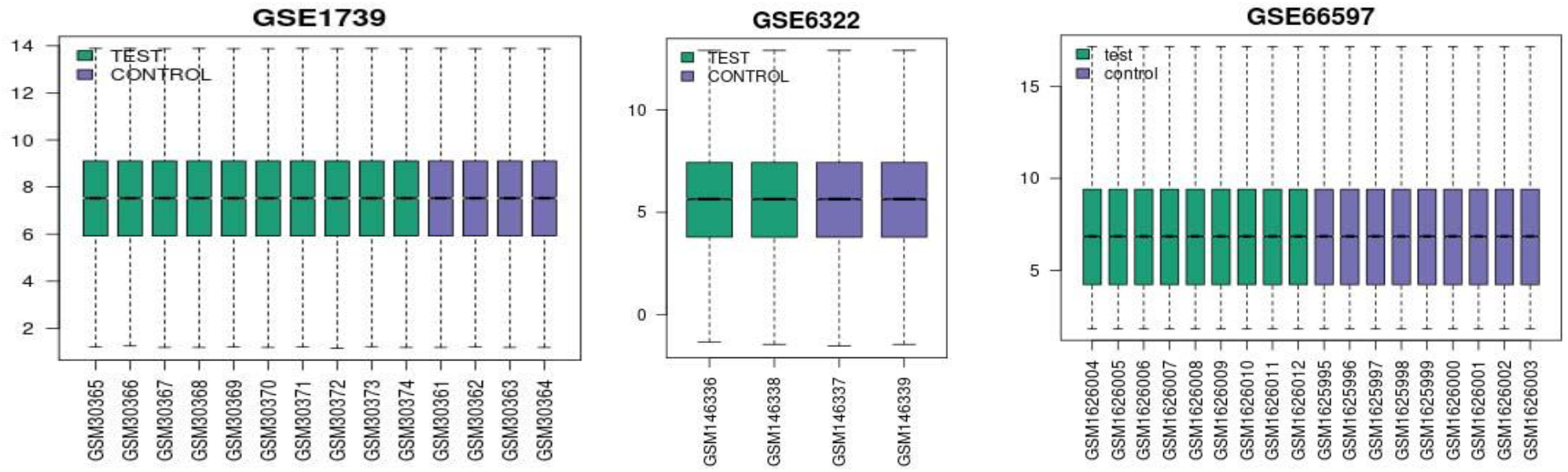
Boxplots showing the differentially expressed genes value distribution of datasets GSE1739, GSE6322, and GSE66597.

### 3.3. Identification of common genes

On the upload of the DEGs to the FunRich online software,50 common genes were identified between dataset GSE1739(SARS-COV) and GSE6322(IAV). However, these 50 common genes are absent in dataset GSE66597(neutropenia) which returned 7 common genes with GSE1739, and 20 common genes with GSE6322 respectively. The diagrammatic representation is shown on the Venn diagram in figure 2.

**Figure 2.**
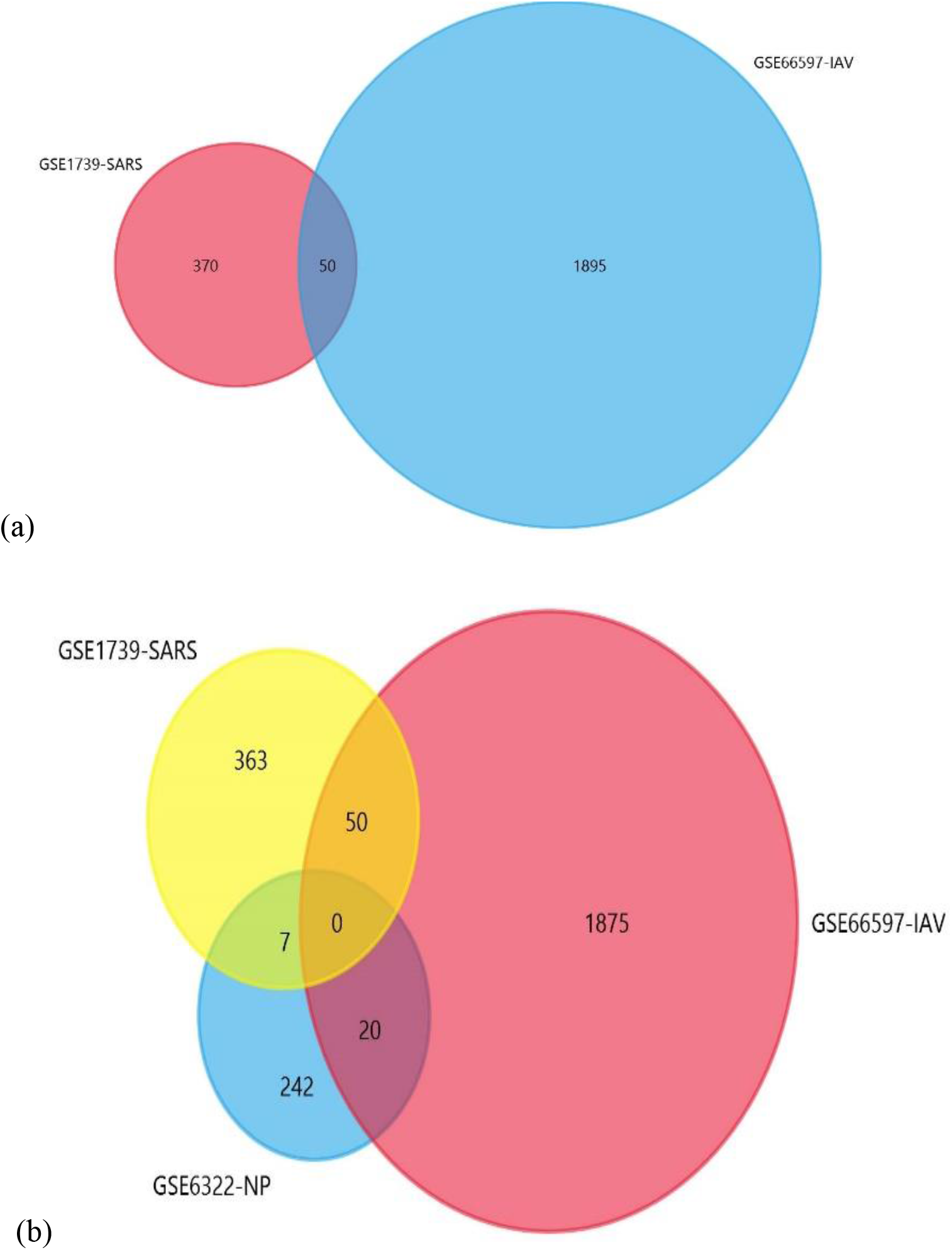
(a) Venn diagram of the DEGs of GSE1739 and GSE66597. (b) Venn diagram of DEGs of GSE1739, GSE66597, and GSE6322.

### 3.4. PPI network of common genes

The diagrammatic representation of the outcome of protein-protein interaction is shown in figure 3. The outcome reveals 50 nodes and 39 edges after concealing disconnected nodes in the network.

**Figure 3.**
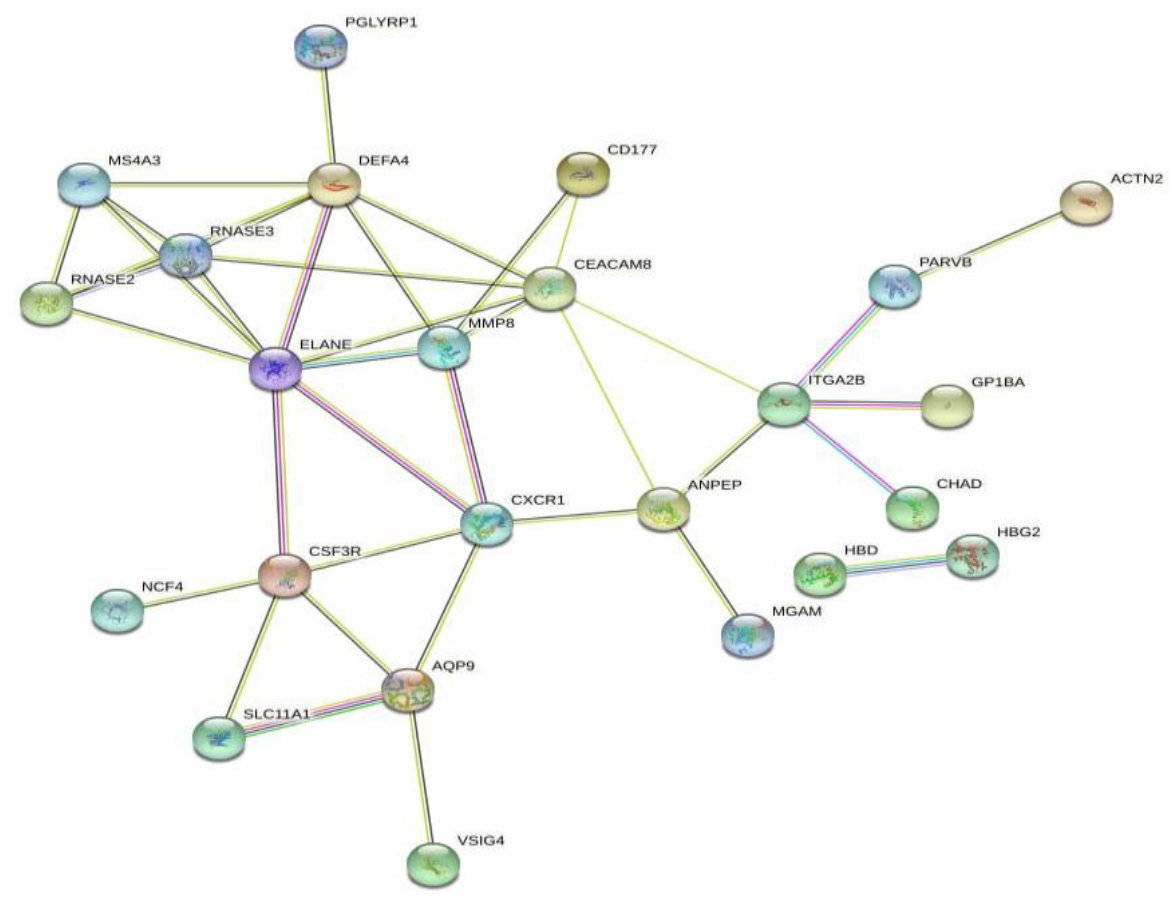
protein-protein interaction (PPI) network constructed by STRING tool.

### 3.5. Identification of hub genes and protein interaction source

After the import of PPI network into the cytoscape desktop tool, the CytoHubba plug-in analysis returned ELANE, DEFA4, RNASE3, RNASE2, MS4A3, CEACAM8, MMP8, CXCR1, CSF3R, ITGA2, regarded as the hub genes. The source of the interactions is revealed using the style function of the cytoscape tool depicted by the direction of arrow has shown in figure 4. ELANE possesses the highest degree of interaction as well as serving as the source of all its interactions.

**Figure 4.**
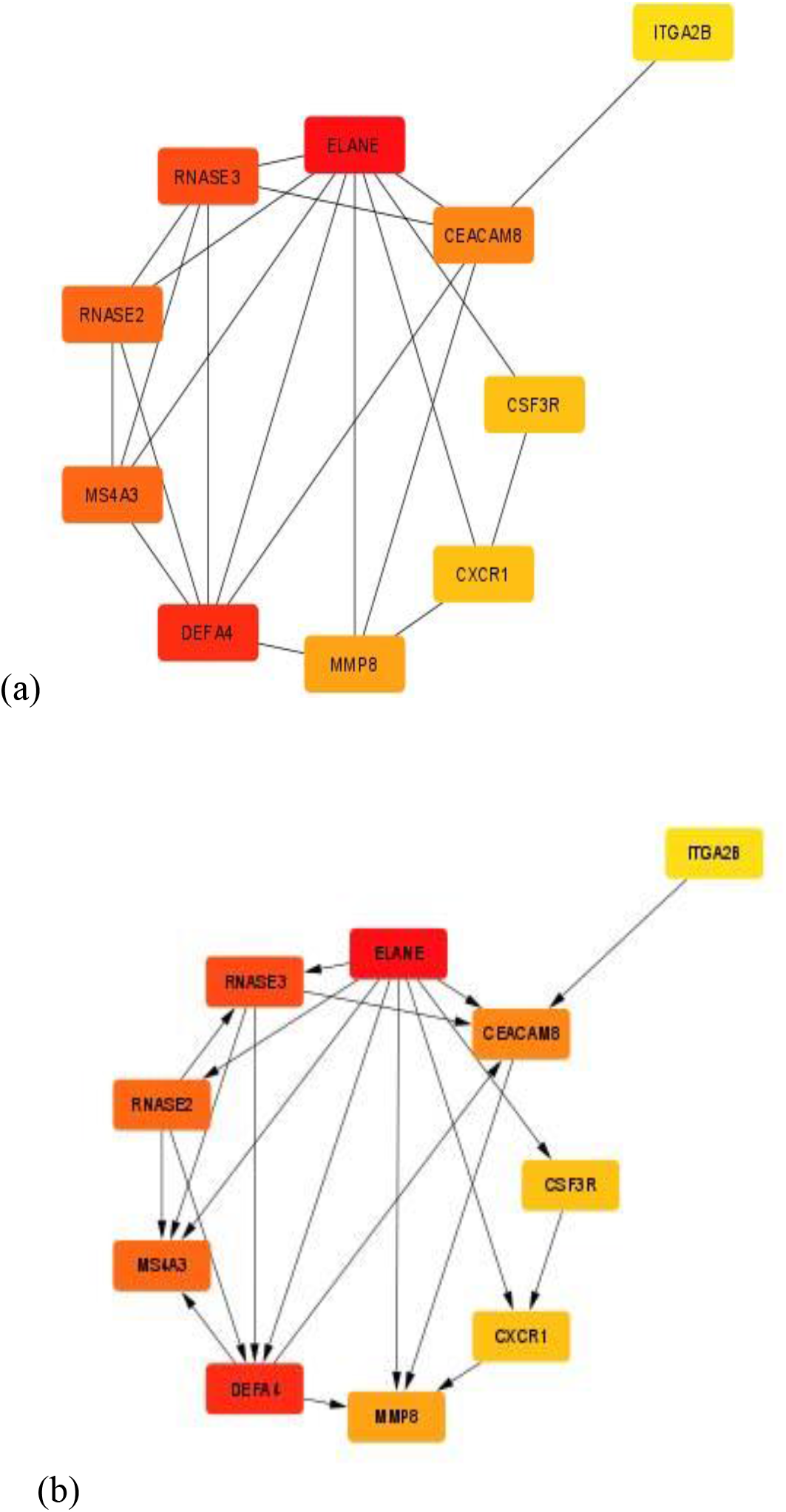
(a) top 10 hub genes. (b) top 10 hub genes with arrows showing direction of interaction. The color represents the degree of connectivity; red color represents the highest degree, orange color represents intermediate degree, while the yellow represents the lowest degree.

### 3.6. Functional and pathway enrichment analysis of hub genes

After the submission of the hub genes in the Enrichr, Enrichr provided several options like GO-biological process (GO-BP), GO molecular function (GO-MF), GO-cellular component (GO-CC). GO analysis showed that the hub genes were significantly enriched in GO-BP such as neutrophil degranulation, neutrophil activation involved in immune response, neutrophil mediated immunity, defense response to gram-negative bacteria, innate immune response in mucosa, mucosal immune response, defense response to bacterium. GO-MF analysis showed that hub genes were significantly enriched in nuclease activity, ribonuclease activity, cytokine receptor activity, serine-type endopeptidase activity, serine-type peptidase activity. For GO-CF analysis, the common genes are significantly enriched with azurophil granules, specific granules, secretory granule lumen, specific granule lumen, secretory granule membrane. The KEGG pathway analysis revealed that candidate genes were highly associated with hematopoietic cell lineage, neutrophil extracellular trap formation, transcriptional mis-regulation in cancer, cytokine-cytokine receptor interaction.

**Figure 5.**
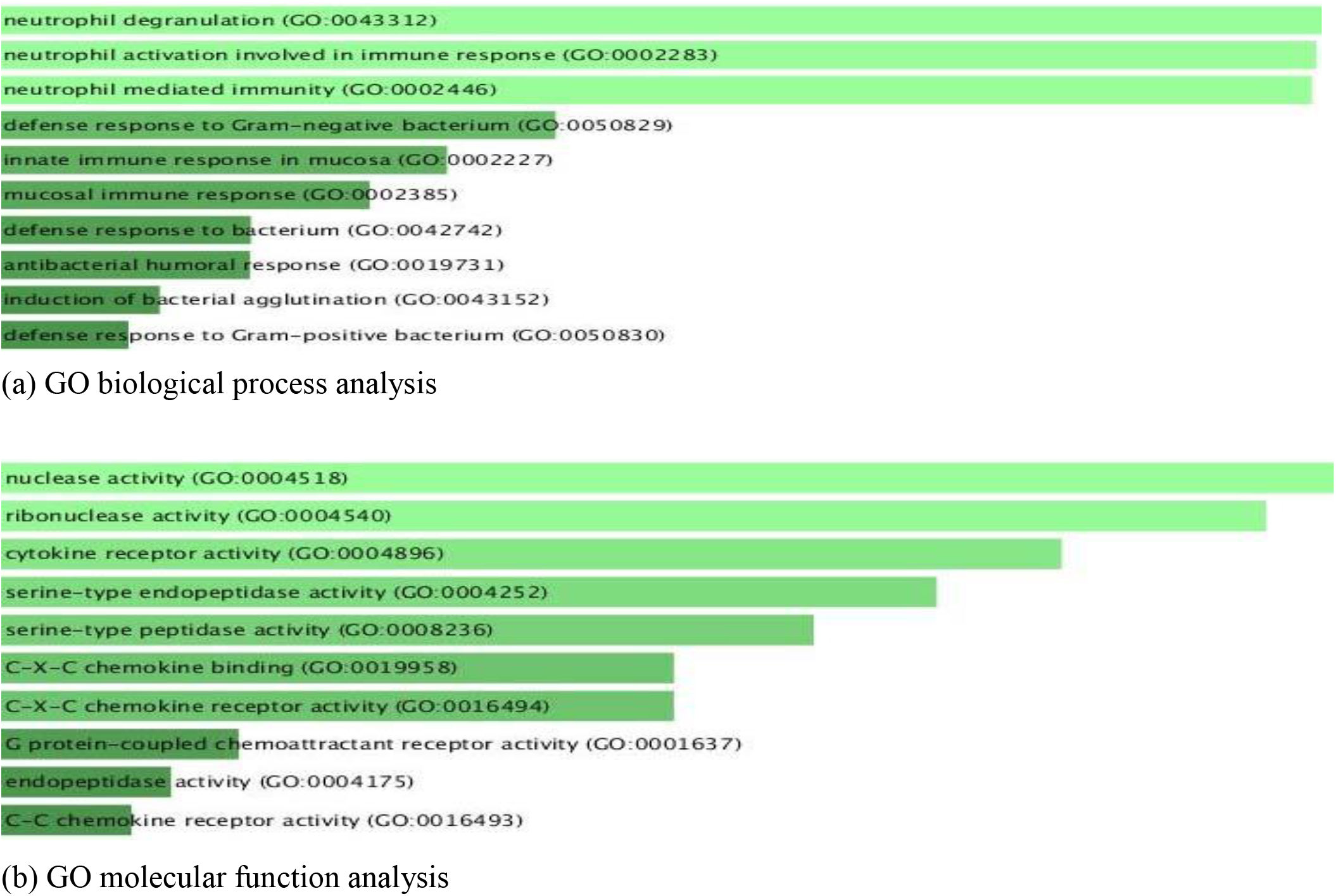
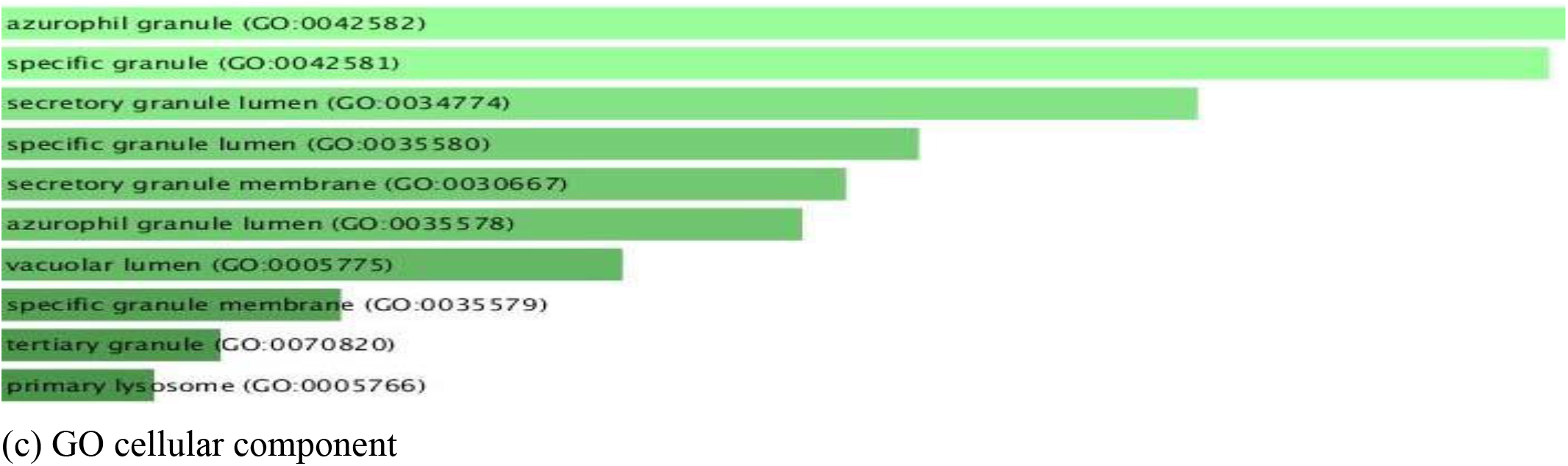
Enriched gene ontology (GO) functions of hub genes.

**Figure 6.**
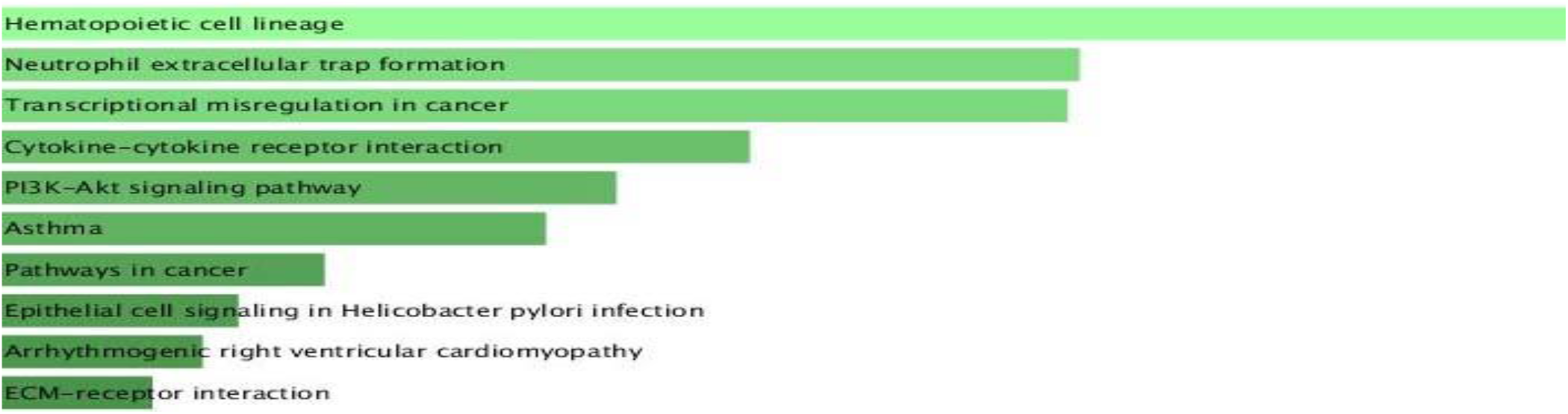
Kyoto Encyclopedia of Genes and Genomes (KEGG) pathway analysis of hub genes.

### 3.7. microRNA interaction analysis

Uploading the hub genes into the online software generated a network between the genes and miRNAs. 14 transcription factors (TFs) and 20 miRNAs were found associated with the hub genes. ELANE has the highest connectivity, with degree score of 10. ITGA2B, CSF3R, SPI1, MS4A3, MMP8, CEACAM8, RNASE3, and DEFA4 possess the degree score of 9,8,6,5,4,3,3,3,2 respectively.

**Figure 7.**
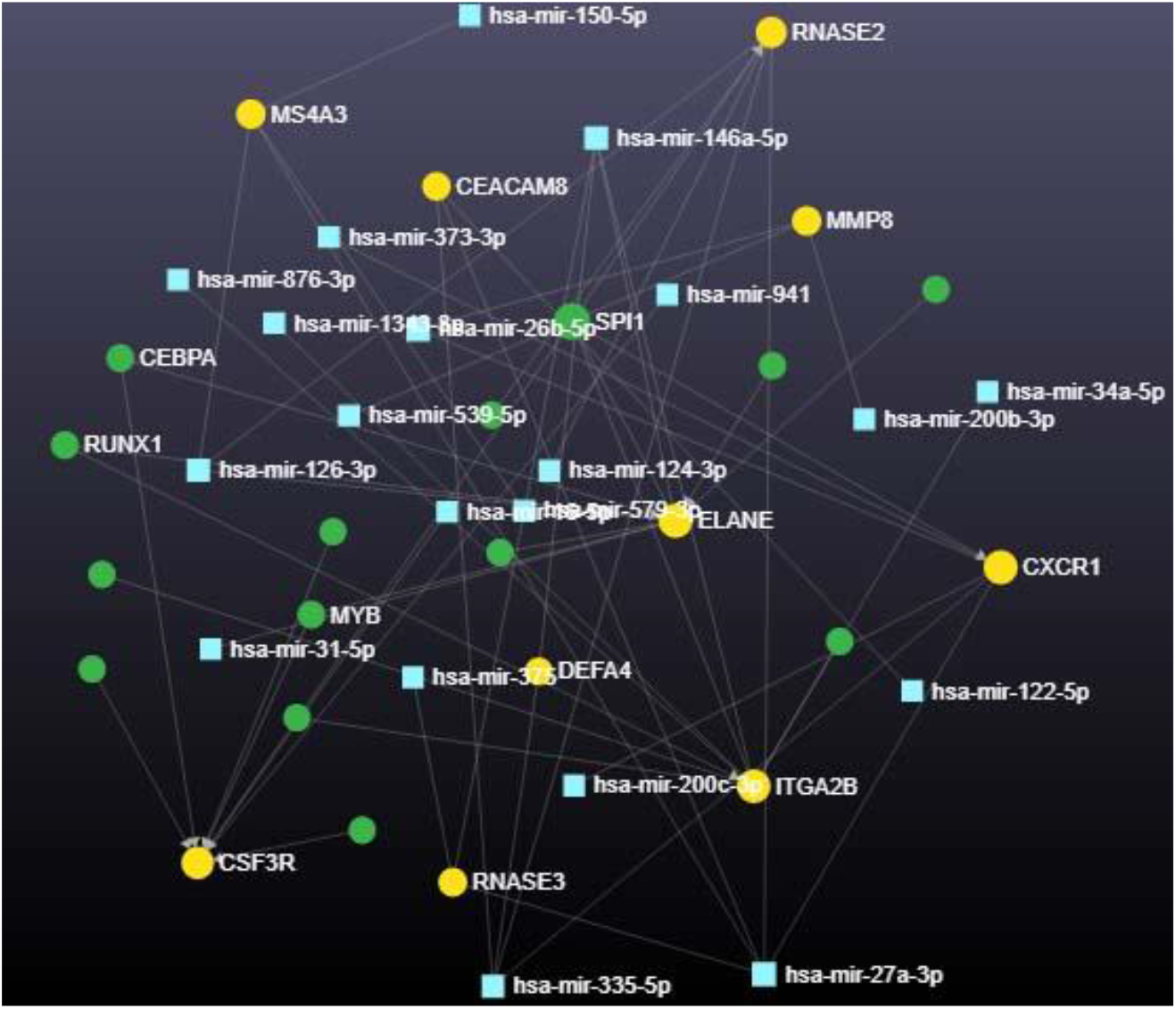
Hub genes and miRNA interaction graph. The genes, miRNAs, and transcription factor TF are represented in yellow, blue, and green colors respectively.

## 4. Discussion

This study evaluated the common genes between SARS-CoV-1 and IAV diseases via a series of bioinformatics tools. After performing the several analyses under set conditions as stated in the methods, we identified some key genes otherwise known has the hub genes which were not previously reported to be common between both diseases, and they include ELANE, ITGA2B, CXCR1, CSF3R, SPI1, MS4A3, MMP8, CEACAM8, RNASE3, and DEFA4. Further evaluation of these genes in respect to NP reveals the deficiency of these identified hub genes in the significantly expressed genes of NP.

Hub genes were investigated in GO term enrichment analysis and KEGG pathway analysis for functional annotation with GO term enrichment analysis, revealing hub genes potential critical roles in defense/immune response through neutrophil degranulation, neutrophil activation, nuclease activity, ribonuclease activity, cytokine receptor activity, chemokine binding, azurophil granules release, specific granules amongst others. Meanwhile, KEGG pathway analysis revealed that hub genes were mainly enriched in hematopoietic cell lineage, neutrophil extracellular trap formation, transcriptional mis-regulation, cytokine-cytokine receptor interaction, P13k-Akt signaling pathway.

Moreover, the characteristic behavior of, Elastase neutrophil expressed (ELANE) gene in relation to the rest of the hub genes makes it a gene of special interest in this work, ELANE gene exhibits a high degree of centrality; which connotes having most interactions in the protein network, suggesting its essentiality in relation to the rest of the obtained hub genes derived from SARS-COV-1 and IAV diseases, as the number of interactions that a protein has in a protein-protein interaction is observed to be correlated with its indispensability [17]. Furthermore, the result shown by figure 4(b) reveals ELANE as the source of signal-flow between its interacting hub genes has represented by the direction of arrows in the diagram, which could possibly suggest its more regulatory function for the ELANE gene in immune response through signaling[18]. This claim might also be corroborated by the significant absence of ELANE and the rest of the hub genes from the DEGs of neutropenia. Neutropenia, has reported in previous reports, is developed by point mutation in the ELANE gene, causing neutrophil the inability to mature with consequent inability to synthesize elastase; a protease protein, which therefore result into smaller-than-normal amount of neutrophil in the blood of the patient, hence susceptibility to infections due to the loss of immune response supposedly to be triggered by neutrophil [11],which makes a lot of biological sense due to the fact that neutrophil respond first at the site of infection/injury, which is then followed by the recruitment of corresponding immune cells and proteins required at the site of infection [19].

## 5. Conclusions

This study has helped us to identify the immune response genes common in both SARS-COV-1 and IAV diseases which has never been reported before in literature. Effort was made to validate the role of these identified genes in immune response by comparing them with DEGs of NP. Moreover, this study has furtherly identified the possible regulatory function of ELANE genes in immune response during viral infection, considering the unique characteristic of this gene in relation to other coexpressed genes. Hence, it is proposed that further search light should be beamed on the mode of interactions and in-vivo studies on signaling pathways of these identified genes, as this insight might come handy in the development of potent inhibitors in the arrest of exaggerated immune response which might portend a negative effect in the form autoimmunity, or in the development of an agonist capable of immunomodulation.

## Acknowledgments (All sources of funding of the study must be disclosed)

This research has no acknowledgement

